# Optical alignment device for two-photon microscopy

**DOI:** 10.1101/312207

**Authors:** Gregorio L. Galiñanes, Paul J. Marchand, Raphaёl Turcotte, Sebastien Pellat, Na Ji, Daniel Huber

## Abstract

Two-photon excitation fluorescence microscopy has revolutionized our understanding of brain structure and function through the high resolution and large penetration depth it offers. Investigating neural structures *in vivo* requires gaining optical access to the brain, which is typically achieved by replacing a part of the skull with one or several layers of cover glass windows. To compensate for the spherical aberrations caused by the presence of these layers of glass, collar-correction objectives are typically used. However, the efficiency of this correction has been shown to depend significantly on the tilt angle between the glass window surface and the optical axis of the imaging system. Here we expand these observations and characterize the effect of the tilt angle on the collected fluorescence signal with thicker windows (double cover glass) and compare these results with an objective devoid of collar-correction. Finally, we present a simple optical alignment device designed to rapidly minimize the tilt angle *in vivo* and align the optical axis of the microscope perpendicularly to the glass window, thereby significantly improving the imaging quality. The performance of the device is demonstrated in an *in vivo* setting and a tilt-correction procedure is described, enabling the accurate alignment (<0.2°) of the cover glass with the imaging plane in only few iterations.

## 1. Introduction

Since its description for biological applications [1], two-photon excitation fluorescence microscopy (TPEF) has become a widespread tool for biomedical research. Compared to other microscopy techniques, TPEF provides key advantages facilitating anatomic and physiologic studies of living organisms: increased imaging depth and reduced phototoxicity [2]. Examples of *in vivo* two-photon microscopy applications range from the investigation of cell structure, dynamics and mobility [3, 4, 5] to fluid transport and blood flow [6, 7].

One of the most common applications of TPEF microscopy is the study of brain cells *in vivo*, which has led to a better understanding of neurobiological processes ranging from the level of single channel dynamics to the large scale functional organization of cortical circuits [8]. Indeed, TPEF provides a high spatial resolution and three-dimensional imaging capabilities, allowing the structural study of neurons and and their subcellular compartments (e.g. dendrites, boutons, spines). Optical access to the brain is typically granted through a craniotomy where the bone is replaced with a glass window. This method allows stable longterm imaging of the same area up to several months, which is essential for elucidating anatomical and functional changes induced by learning or development [9].

Cranial windows are manufactured from one [9] or more layers [10, 11] of coverslip glass typically yielding a total glass thickness of 100 to 400 μm. Multiple layers of glass can help prevent bone regrowth and provide additional mechanical stability. However, the optical aberrations induced by the refraction index mismatches between the brain tissue and the immersion medium [12] are further aggravated by the refractive index and the thickness of the glass window. As a result, the signal intensity and optical sectioning capabilities are reduced, thereby decreasing the quality of the images and the attainable depth [13, 14]. Adaptive optics can optimally recover imaging quality by correcting all aberration modes [15], yet when spherical aberrations are dominant, the most accessible approach for improving image quality involves adjusting the correction collar present in many high numerical aperture water immersion objectives [16]. Nevertheless, the theoretical optimal reduction in spherical aberrations induced by the correction collar requires the optical axis of the objective to be perpendicular to the cranial window.

Unlike *ex vivo* preparations, the orientation of the imaging window in awake, head-fixed animals is imposed by the overall body position and the location of the brain area to be imaged. This often leads to an undesired tilt angle between the optical axis of the microscope and the orientation of the glass window. Modern two-photon microscopes used for *in vivo* research are therefore typically equipped with rotating frontends (e.g. https://www.janelia.org/open-science/mimms) allowing for tilt angle correction. As described earlier, proper adjustment of the cranial window orientation relative to the microscope objective is crucial, since tilted windows lead to additional optical aberrations that cannot be compensated for with the correction collar. Moreover, if a tilt angle is present, collar correction can even have a deleterious effect on image quality [17]. A number of strategies have been devised to align the window, such as visual inspection, fluorescent microbeads placed on the window [18] and wavefront sensing [17]. From our experience, visual inspection typically leads to an incomplete tilt correction and is thus suboptimal. Wavefront sensing allows precisely measuring the tilt angle but requires an experienced user and costly equipment. Finally, application of fluorescent microbeads on the cranial window is an interesting strategy but requires placing beads on the window. Technologies for rapid and robust alignment of samples with respect to the optical axis are therefore needed.

Here we build on our previous observations by characterizing the impact of different tilt angles and cover glass thickness on the intensity and axial profiles of fluorescent microbeads and show the impact of a tilt angle on the quality of *in vivo* images. To minimize the tilt during biological experiments, we developed a simple optical device designed to facilitate the alignment of the optical axis of the microscope with respect to the cranial window. Finally, we measured the sensitivity of the device and provide a protocol for rapid and reliable alignment of the optical axis with the glass window.

## 2. Methods

### 2.1 Optical System

#### 2.1.1 Microscope for bead imaging

Fluorescent beads were imaged on a TPEF microscope equipped with adaptive optics [17]. Fluorescence was excited by a pulsed near-infrared laser (920 nm, Coherent, Chameleon Ultra II). The excitation laser was reflected off a deformable mirror (Alpao, DM 97-15) after 10fold expansion by two lenses (focal lengths: 50 mm and 500 mm). The deformable mirror was conjugated to a pair of galvanometer mirrors for point-scanning (Cambridge Technology, 6215H) and to the back focal plane of an objective lens by pairs of lenses (focal lengths: 300 mm and 100 mm between the deformable mirror and the first galvanometer mirror, 85 mm and 85 mm between the two galvanometer mirrors, and 100 mm and 400 mm between the second galvanometer mirror and the objective back focal plane). The high numerical aperture objective lens (Nikon, CFI Apo LWD 25XW, 1.1 NA and 2 mm WD) was used to focus the excitation light onto the sample and collect the emitted fluorescence. The emitted fluorescence was separated from the excitation light by a dichroic mirror (Semrock, FF665-Di02-25×36) positioned close to the objective lens. The fluorescence signal was then focused onto a photomultiplier tube (Hamamatsu, H7422-40) by a lens (focal length: 75 mm) and spectrally filtered by a bandpass filter (Semrock, FF03-525/50-25). Only aberrations intrinsic to the optical microscope were corrected by the deformable mirror such that the microscope would provide a diffraction-limited focal volume when imaging beads in the absence of sample-induced aberrations. A goniometer stage (Thorlabs, GNL10) was used to control the tilt angle of the samples.

#### 2.1.2 Microscope for *in vivo* brain imaging

*In vivo* imaging was performed as previously described [19] using a custom made TPEF microscope with a rotational frontend (MIMMS; https://openwiki.janelia.org/wiki/display/shareddesigns/MIMMS) controlled by ScanImage 5 (https://vidriotechnologies.com). The microscope was equipped with a 25× collar correction objective (Nikon, CFI Apo LWD, 25XW, 1.1 NA and 2 mm WD) and a resonant scanner head (Thorlabs). Fluorescence was excited by pulsed near-infrared laser (940 nm, Coherent, Chameleon Ultra II) corrected with a group velocity dispersion compressor (Coherent, Chameleon PreComp). The laser power was modulated with a Pockels cell (Conoptics, 35080-LA-02) to a maximum of 15 mW (measured in air at the focal plane). 512×512 pixels images were acquired at 29.38 frames/s using a gated photo multiplier tube (Hamamatsu, H11706P-40 SEL) and digitally written in 16-bit format to disk. The angle of the microscope head was measured using a digital angle gauge (Wixey).

### 2.2 Bead imaging and analysis

Fluorescent beads (Invitrogen, FluoSphere carboxylate-modified microsphere, yellow-green 505/515, 2 μm in diameter) were placed on a microscope slide, on which a thin layer of polylysine had been applied and immobilized these beads via electrostatic force. A drop of water was put directly on the beads before adding any cover glass (Fisher Scientific, No. 1.5, 160-190 μm thick) to ensure no additional wavefront aberration would be introduced by air gaps; the cover glass mimicked a cranial window and was the only source of aberration. The sample was placed on the goniometer stage and its tilt angle was adjusted as needed. Image stacks were recorded for isolated beads at an axial step size of 0.2 μm to capture their three-dimension fluorescence distribution. To assess the effect of sample tilt as a function of the cover glass thickness, images were acquired in a single or double cover glass configuration at 5 tilt-angles: θ= 0, 1, 2, 5, and 10° degrees. This was repeated twice: once with the objective correction collar adjusted to zero and not correcting for any glass thickness (“no correction”) and again with the objective correction collar adjusted to the thickness of the glass window (“collar correction”).

Image analysis consisted of quantifying the axial full width at half maximum (FHWM) and the TPEF intensity for each bead, in order to characterize the effect of the tilt and glass thickness on imaging. The TPEF intensity of green beads was measured using ImageJ. First, the background was subtracted using the “rolling ball” algorithm on these image stacks. A maximal intensity projection along the axial direction was then performed to produce a twodimensional image, on which a Gaussian blur (radius: 0.1 μm) was applied. The bead intensity was measured as the peak intensity within the bead. The intensity was normalized to the value at a tilt angle of θ= 0° and under a single cover glass for Fig. 1A and to the value at a tilt angle of 0° and with collar correction for Fig. 1B. Maximal intensity projections along both lateral axes were used to produce axial profiles on which the axial FWHMs could be evaluated. The FWHM values were estimated from the profiles by finding the two points neighboring half of the maximal intensity value on each side of the central peak and then performing a linear interpolation. At least seven independent bead measurements were performed for each experimental group. Statistical analysis was performed in R using linear regression models.

### 2.3 Animal preparation and imaging

#### 2.3.1 Cranial window surgery on Thy1-GFP line M mice

All experiments were approved by the Animal Care Committee of the University of Geneva and by the Direction générale de la santé of the Canton of Geneva. Mice were held under a controlled 12 hr light/dark cycle (7:00 lights on, 19:00 lights off) with *ad libitum* access to food and water. *In vivo* imaging was performed on one adult female GFP-M mouse (12 months old). GFP-M mice express EGFP under the control of a modified Thy1 promoter (Tg(Thy1-EGFP)MJrs/J) leading to sparse labeling of cortical neurons. Cranial window surgery was performed as previously described [19]. Surgical procedures were conducted under aseptic conditions and isoflurane anesthesia (1.5%) in a custom-made stereotactic apparatus equipped with a thermic plate (37°C). After anesthesia induction, toe- and tail-pinch nociceptive responses were assessed and anti-inflammatory (2.5 mg per kg body weight dexamethasone intramuscular (i.m.); 5 mg/kg Carprofen subcutaneous (s.c.)), analgesic (0.1 mg/kg buprenorphine i.m.) and local anesthetic (1% lidocaine s.c. under the scalp) drugs were administered. The scalp was cleaned with ethanol 70%, disinfected with Chlorhexidine and excised with surgical scissors. The periosteum was removed with corneal scissors and the blunt edge of a scalpel blade. The bone was gently roughened to increase adhesion and a custom-made titanium headbar was glued with cyanoacrylate glue. The head-bar was secured with dental cement and left to cure for a few (~10) minutes. A craniotomy (~2 by 3 mm) was performed with a dental drill equipped with a round bur on the left frontal cortex. The edges of the craniotomy were flattened with a flat-end cylinder bur to improve window-bone fitting. The excised bone was replaced by a matching-shape coverslip window. Two glass windows were manually cut from 150 μm thick coverslips (Menzel-Gläser #1) with a diamond scribe (Fiber Instruments, FO90C). One of the windows approximated the shape and size of the craniotomy in order to replace the excised bone, while the other one was slightly larger in order to sit on the remaining bone edge of the craniotomy. After cleaning with ethanol 70%, both glasses were glued together with optical glue (Norland adhesive 61) and cured with UV light for 1 minute, producing a double layer cranial window. Excess of the optical glue was removed and the cranial window cleaned with ethanol 70%, rinsed with sterile saline. Subsequently, the cleaned window was gently positioned on top of the brain and secured against the bone with cyanoacrylate glue and covered with dental cement. After the surgery, the mouse was returned to its home cage and allowed to recover for several days before experiments.

#### 2.3.2 *In vivo* imaging and analysis

*In vivo* imaging was performed under isoflurane anesthesia (1.5% in oxygen). The body temperature of the mouse was kept at 37°C using an electric heating pad. The head fixation system was mounted on a goniometer with a rotation range of ±10° (Thorlabs, GN1). After alignment of the cranial window with the optical axis (using the tilt correction device described in this paper), stacks of GFP expressing neurons in layer 2/3 of the frontal cortex were acquired at different tilt angles: θ= 0, 5, and 10° degrees. Stacks were performed with a step size of 2 μm in the z axis and a pixel size of 0.4 μm (x and y) for imaging the soma of neurons. For imaging neuronal processes (dendrites and axons), the step size was set to 1 μm and the pixel size to 0.15 μm. Each slice of the stack was obtained by averaging 30 or 50 frames. After having acquired a stack at θ= 0°, the tilt angle was changed using the goniometer and the field of view was manually adjusted at the soma level to match the imaging plane at the center of the field of view. The impact of tilt angles at 0°, 5° and 10° was sequentially assessed by performing 3 independent measurements for each angle, at the same location. Images of neuronal processes were analyzed using ImageJ and custom written MATLAB scripts. Regions of interest containing neuronal processes with boutons were projected in the z axis (maximal intensity, 15 slices equivalent to 15 μm) within an area of 50 by 512 pixels (7.3 × 75 μm) and the intensity profile was obtained. Intensity profiles of different acquisitions were aligned using a peak alignment algorithm *(msalign* function, MATLAB).

### 2.4 Tilt alignment device

#### 2.4.1 Construction and calibration of the device

The alignment system is a small (12 × 6 × 3 cm) optical device rotating around the TPEF microscope objective. It allows determining and correcting the tilt of the cranial imaging window in relation to the optical axis of the microscope. It was designed combining both readily available commercial parts and 3D printable elements designed in Fusion 360 (Autodesk). A detailed part list and access to the 3D blueprints are available at https://huberlab.org/resources/optical-alignment-device. 3D pieces were printed in polylactic acid (PLA) with an Ultimaker 3 printer. A miniature laser diode (Roithner Lasertechnik, APCD-650-01-C3) was lodged into an articulated arm composed of a swivel coupler (Thorlabs, C2A), a shortened rod (Thorlabs, ER90C) and a 3D printed adapter. The size of the laser beam was decreased by passing through a 200 μm wide pinhole (Thorlabs, P200H) mounted in front of the diode. The laser diode was controlled by a micro-switch (Distrelec, 110-34-092) and powered by a lithium 3V battery (CR 1/2 AA, Distrelec, 169-01-539), which was fixed to the device using a battery holder (Distrelec, 169-52-048). Opposite to the laser, a millimeter graded paper was glued on a 3D printed target to determine the displacement of the laser beam reflected by the cranial window.

First, to illustrate the physical principle of the device, we characterized its performance under well-controlled automatized settings. A cover glass fixed on top of a couple of orthogonally aligned goniometers was used to emulate different tilt angles (θ) of the cranial window. To facilitate the measurements and avoid any potential movement artefacts, the device position was kept fixed while the sample was rotated instead. As such, the goniometer-cover glass ensemble was mounted on a rotating motorized stage (Zaber, X-RSW60C-E03, controlled with a custom MATLAB script), which was used to emulate the rotation angle (γ) of the device. The motion of the reflected laser beam on the target was captured during the rotation of the motorized stage using a Firefly MV FMVU-03MTM camera (Pointgrey) and the MATLAB Image Acquisition toolbox. Next, we assessed the performance of the device in real-experimental *in vivo* settings using n = 3 mice implanted with cranial windows in the frontal cortex. Mice were anesthetized with isoflurane and head-fixed on a rotating platform (Thorlabs breadboard). After the cranial window tilt was minimized using the protocol described herein (see 3.2.3), a miniature Rasperry Pi Spy Camera (Adafruit, 1937) was fixed on the target to capture the laser beam displacement at different rotation angles of the device (γ) and determine the residual tilt of the cranial window. For each of the procedures described above, extraction of the beam spot and its centroid position for each rotation angle γ was performed offline in MATLAB.

## 3 Results

### 3.1 Image brightness and resolution depends on cover glass orientation angle and collar correction

To increase imaging stability during TPEF imaging in behaving animals, it is customary to prepare cranial windows with two or more layers of cover glass glued together with optical adhesive (e.g. [10, 11]). We have previously demonstrated the deleterious effect of sample tilt on imaging quality under a cranial window made of a single layer of cover glass [17]. Considering that the increased glass thickness in double-layered windows should introduce even larger optical aberrations, we investigated the effect of sample tilt and glass thickness on the images of fluorescent microbeads (2.0 μm in diameter). Images were acquired at tilt angles of θ= 0°, 1°, 2°, 5°, and 10° behind single- or double-layered windows, with the collar adjusted to the position corresponding to the window thickness (Fig. 1A). The size of the fluorescent microbead was chosen so as to mimic real neuronal structures, such as spines.

As expected, we found that collar adjustment provided optimal signal recovery at zero tilt angle, whereas increasing the tilt angle significantly decreased the bead intensity (p-value: 2×10^−16^, Fig. 1A). For example, in the absence of tilt, spherical aberrations were reduced by the collar correction, and beads under a double-layered window had a similar intensity to those under a single-layered window. Signal deterioration with tilt angle was steeper for double-layered windows than for single-layered windows (p-value: 2 × 10^−16^, Fig. 1A): a tilt angle of only θ= 2° reduced the intensity by 35% for beads under a single-layered window but by 70% for beads under a double-layered window. Increasing the tilt angle θ further led to larger aberrations (e.g., coma [17]) for double-layered windows, and larger decreases in fluorescence intensity.

As the correction collar compensates for spherical aberrations, its efficiency is optimal solely when the cover glass plane is orthogonal to the optical axis. As such, the improvement in image brightness with and without collar correction is maximal at minimal tilt angle (Fig. 1B). Here, under a single-layered window with increasing tilt angle θ, the signal recovery after collar correction rapidly deteriorated, with maximal signal recovery (~ 10× increase) observed at zero tilt.

Comparisons of the axial FWHM profiles of beads under single- and double-layered windows at different tilt angles confirm the same trends (Fig. 1C, D). Only at θ= 0° and 1° tilt angles, collar correction yielded axial FWHMs close to the diffraction-limited value. The axial FWHMs of the beads increased progressively with tilt angles and at the same tilt angles, double-layered windows always led to larger axial FWHMs (p-value: 2 × 10^−16^), corresponding to the worsening of axial resolution. Interestingly, at larger angles (e.g., θ= 10°), collar correction led to a worse axial resolution than in an uncorrected configuration (cf., axial FWHMs at θ= 10° in Fig. 1C and Fig. 1D). Nevertheless, tilt angle and glass window thickness do not affect the lateral FWHM profiles of beads [17]. To further confirm the deleterious effect of a tilt in the cranial window under *in vivo* conditions, we qualitatively compared structural images of cortical neurons obtained at different imposed tilt angles (Fig. 1 E, F). We imaged cell bodies and cortical processes in a mouse line displaying a sparse population of green fluorescent protein expressing neurons (Thy1-GFP line M). As highlighted by tiles E and F of Fig. 1, the image quality quickly deteriorated with increasing tilt angles θ, confirming the detrimental effects of such misalignments under experimental imaging conditions. This effect was particularly evident for small structures such as boutons, spines and axons (Fig. 1F), but also affected the intensity of larger structures such as the soma of neurons.

Taken together, these results indicate that image quality rapidly degrades when the cranial window is tilted in relation to the imaging plane of the microscope and that multiple layers of glass worsen this effect. Moreover, adjusting the objective correction collar alone is not sufficient to recover image quality. Ultimately, diffraction-limited imaging performance can only be recovered by both minimizing window tilt and adjusting the objective correction collar.

### 3.2 Tilt correction device

The results presented above highlight the effect of the tilt angle θ of the cover glass with respect to the optical axis in in vivo imaging. Minimizing this offset angle is therefore critical to ensure optimal imaging performance, as it can lead to deleterious aberrations and strongly reduce the signal-to-noise ratio as well as achievable imaging depth. Although several strategies have been devised to align the cover glass parallel to the imaging plane, these strategies can be imprecise, time-consuming or costly. The device presented below addresses these issues as it is based on commercially available components and printable parts. It enables estimating and minimizing the tilt angle θ of the sample (i.e. animal’s cranial window) by adequately adjusting the microscope frontend (β) and the animal’s (α) rotation angle (Fig. 2A). For clarity, the following terms are depicted in Fig. 2A-B: the microscope plane is the plane containing the optical axis and within which the frontend of the microscope can be rotated; the table plane corresponds to the plane in which the head-fixed animal can be rotated; γ is the azimuthal rotation angle of the device.

#### 3.2.1 Operational principle

The device presented in Fig. 2A allows an accurate estimation of the angle θ between a reflective surface (i.e. cranial window cover glass) and a reference plane (imaging plane) by assessing the changes in the reflectance angle of a laser beam impinging on the cover glass at varying rotational angles γ (azimuth angle around the objective) and at a given incidence angle φ. By placing a target (with scaled gradations, i.e. millimeter paper) on the opposite size of the laser source, the reflectance angle is converted into a position. The tilt angle θ of the glass surface can be inferred as an amplitude difference by measuring the position of the laser spot on the target at two opposing azimuth angles (i.e. γ= 0° and 180°). In the first case, depicted in the top panel of Fig. 2C, the reflection angle of the beam is given by: φ’_0°_ = φ − 2θ. In the opposite case, shown in the bottom panel of Fig. 2C, the angle is obtained as φ’_180°_ = φ+ 2θ. Consequently, the difference between both angles is independent of the incidence angle φ, and can be calculated as Δφ = φ’_180°_ − φ’_0°_ = 4θ (Fig. 2D). From this observation, and as shown in Fig. 2D, the relation between this amplitude and the distance between the incidence positions of the beam on the surface and the target L and the surface tilt angle θ can be derived by the following Equation 1:

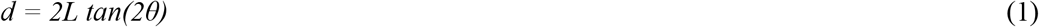

The rotation angle α of the animal is determined by finding the rotation angle γ of the device displaying the maximal or minimal position of the laser spot on the target.

**Figure 1.**
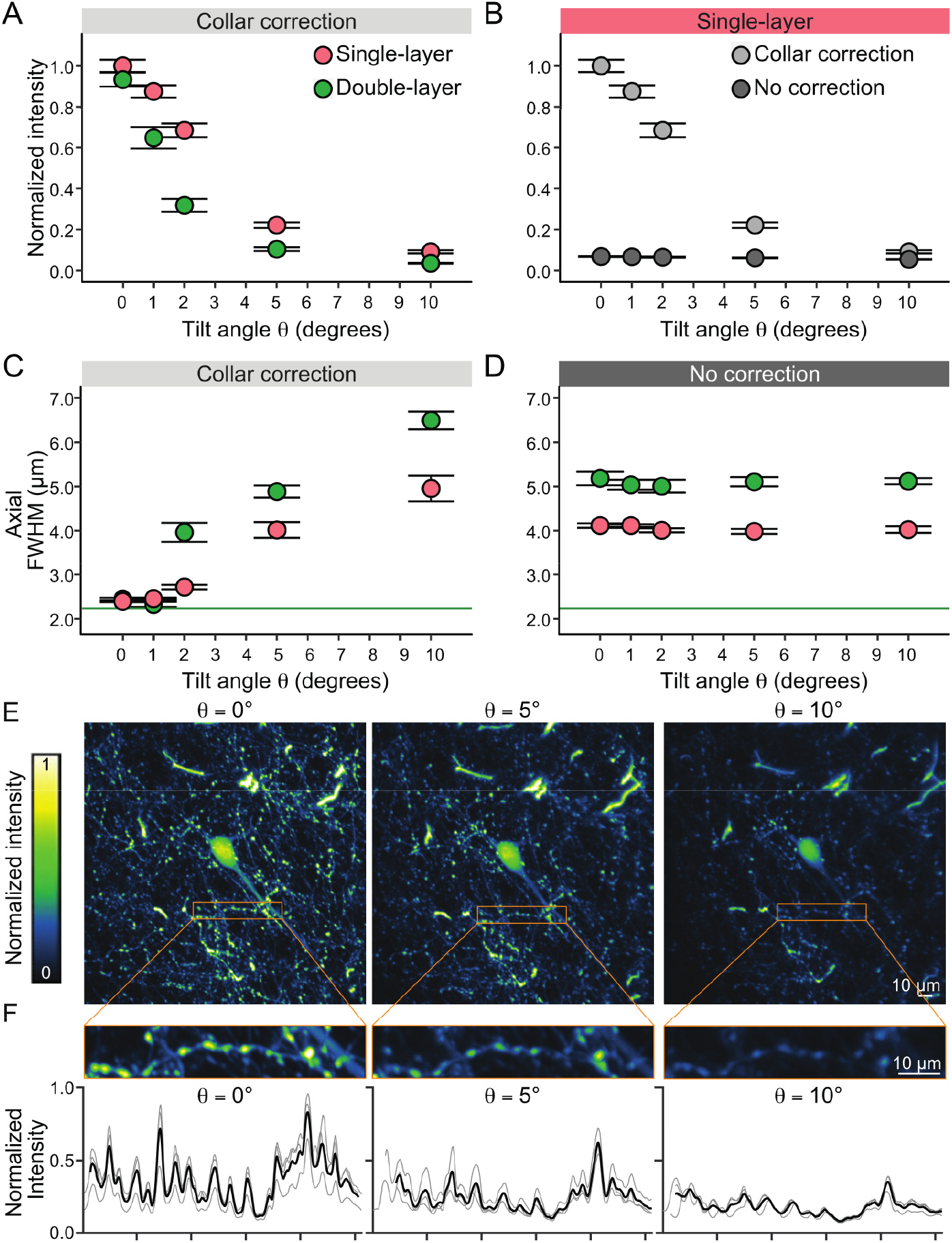
Two-photon fluorescence intensity and FWHM profiles of fluorescent microbeads under different orientation angles, cover glass layers and collar-corrections. (A) Increasing the tilt angle between the optical axis of the two-photon microscope and the axis perpendicular to the imaging window significantly decreases light intensity measured from individual microbeads with a collar corrected objective. Even at small tilt angles of θ = 1° a significant loss of intensity is detected. This effect is amplified when the thickness of the cover glass is doubled. (B) The lack of collar correction drastically decreases the measured bead intensity. (C) With a collar correction, the axial FWHM is fully corrected at θ = 0° tilt angle close to the theoretical limit (green solid line) for single and double cover glass preparations. Increasing the tilt angle significantly degrades the axial resolution. (D) Without collar correction the axial resolution is deceased. The double glass configuration further degrades the axial resolution. The resolution of the uncorrected condition is comparable to a θ = 5° tilt in the corrected condition. (E) Z projection (maximal intensity) of 7 slices (14 μm) at the soma level of a layer 2/3 neuron imaged in the frontal cortex at a 5x magnification factor. The same neuron was imaged at different tilt angles (θ = 0°, 5°, and 10°). (F) Top: zoom in of a neuronal process with boutons acquired at 15x magnification factor. The maximal intensity Z projection (15 slices, 15 μm) was used to obtain the intensity profile of the region of interest on 3 different acquisitions at each tilt angle. Bottom: intensity profiles were aligned to the peaks (thin traces) and averaged (thick traces) for comparison across conditions. Scalebar: 10μm

#### 3.2.2 Device characterization

We first validated the model described in Eq. 1 by placing a cover glass on a 2-axis goniometer mounted on a motorized rotational stage and positioned under the microscope. The alignment device was placed on the microscope’s objective and the laser beam was directed towards the rotating cover glass. The pitch and yaw of the laser diode were adjusted to ensure that the spot was at the center of the rotating sample and that its projection could be read on the target for each rotation angle γ of the motorized stage. The position of the spot on the target was recorded for different rotational angles γ and 5 different tilt angles of one of the goniometers from θ= 0° to 4°. The θ= 0° angle of both goniometers were found by minimizing the amplitude of the beam on the target (only one of the goniometers was tilted for these experiments, the other goniometer was maintained at its “flat” angle). As shown in Fig. 3A and B, as the device is rotated around the objective, the spot wanders on the target, creating an ellipsoid with the amplitude varying with the sample’s tilt angle θ. As expected, when the cover glass is flat (i.e. θ= 0°), the laser beam is reflected at the same spot in the target regardless of the device’s angle γ. We repeated this measurement at different tilt angles over 10 trials and report the results on Fig. 3C showing a good agreement between the geometrical model (bold line, Eq. 1) and the experimental measurements (red circles).

**Figure 2.**
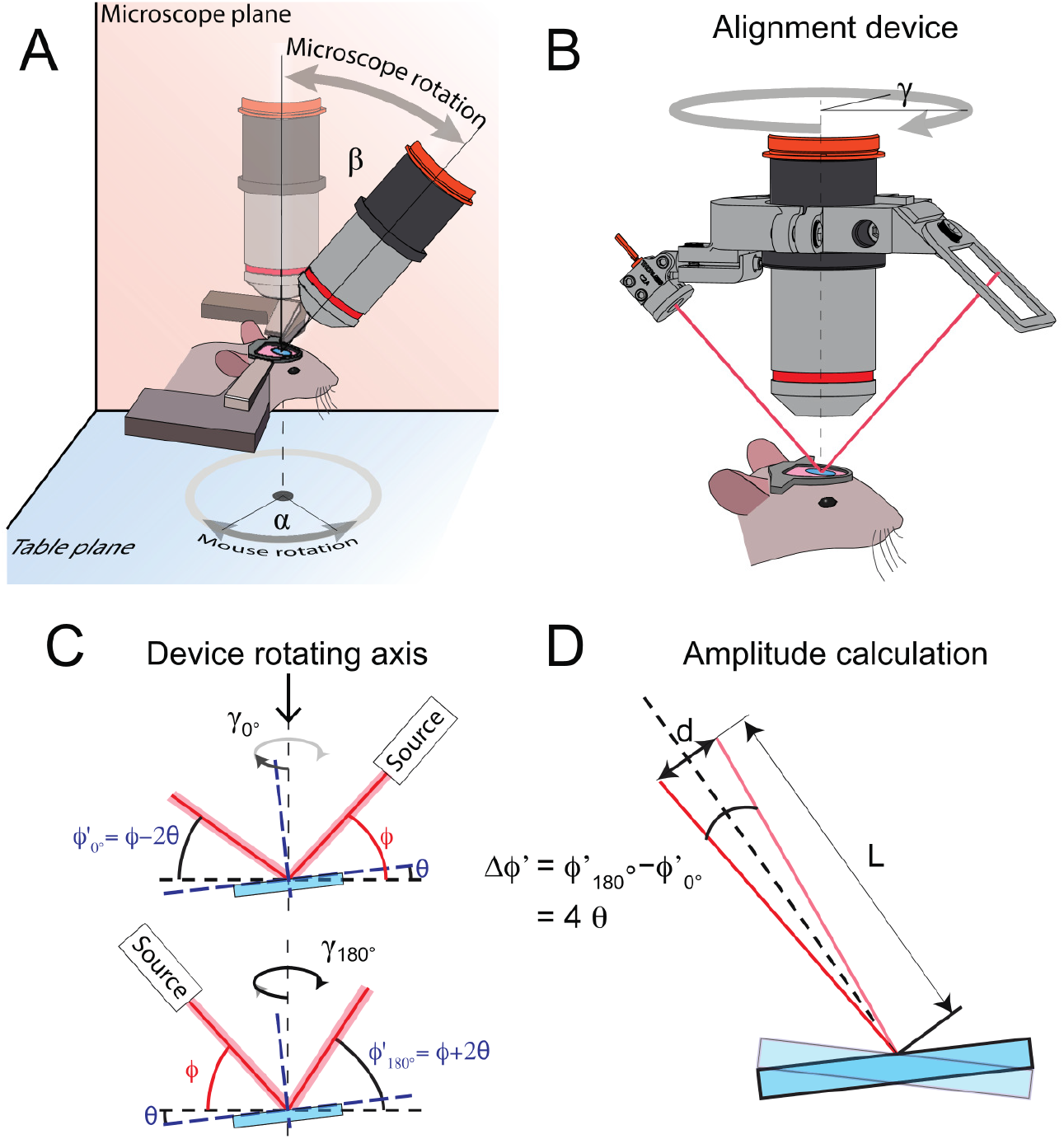
Tilt angle measurement principle. (A) In our configuration, the microscope frontend can be rotated along the microscope plane (in orange) by an angle β and the mouse around the table plane (in blue) by an angle α. (B) The tilt angle can be estimated by mounting the device on an objective. The device comprises a laser source and a target screen placed on opposite sides of the rotating ring. (C) If a tilt θ is present, the reflection angle of the laser beam (red line) will vary depending on the orientation of the device around the optical axis (dotted line), as shown in this panel. (D) This variation in reflective angle φ’ will entail a change in position on the target d, depending on the tilt angle θ and the distance L between the cover glass window (blue) and the target.

#### 3.2.3 Tilt adjustment protocol

Once the alignment-device is placed on the objective (or objective holder) and the animal is set under the microscope, the following steps will enable adjusting the microscope head (or conversely the animal holder if the microscope does not enable rotating the objective) to minimize the angle between the imaging plane and the tilt angle θ of the first surface of the glass window. For the following steps, we define the zero of the angle of the alignment-device γ= 0° when the plane comprising the laser diode beam is parallel to the microscope plane (Fig. 2A). The microscope head tilt β is defined as β= 0° when the optical axis of the objective is perpendicular to the table plane (Fig. 2A). Finally, the tilt angle θ of the cover glass is defined as θ= 0° when it is parallel to the table plane.

1. The laser diode is turned on and the position of the animal is adjusted on the table plane so that the laser beam is reflected by the glass surface of the cranial window and hits the target screen. The frontend of the microscope should be adjusted so that β= 0°(i.e. the objective is straight).
2. To facilitate the tilt estimation, the transverse motion of the laser spot on the cranial window during the rotation of the alignment-device should be minimized (to obtain a continuous reading of the spot on the target). This can be obtained by adjusting the distance between the objective and the head of the animal. If the spot rotates in the same direction as the alignment-device, the distance between the objective and the animal has to be increased, and vice-versa. The distance between the laser spot on the glass window and its projection on the target is measured, for example with a calibre (this will provide the value of L in Eq. 1).
3. The rotation angle of the animal (α) is adjusted as follows: the alignment-device is rotated around the objective and the spot movement is observed on the target. If the cranial window is misaligned with respect to the optical axis of the microscope, the laser will be reflected on the target forming an ellipse (as shown in Fig. 3B). Next, the rotation angle of the alignment-device (γ_1_) at which the spot is at the maximum or minimum position of the ellipse is determined. The animal (α) is rotated on the table plane by γ1 degrees to ensure that the maximum and minimum longitudinal position of the ellipse is aligned with the microscope plane (i.e. parallel to the orange plane in Fig. 2A).
4. To estimate the tilt θ, the alignment-device is rotated around the objective and the amplitude d of the projected ellipse is measured along the longitudinal axis of the target (i.e. the difference in height between the maximum and minimal position). This distance is converted into an angle θ1 using Eq. 1.
5. The microscope rotation angle β is moved using the tilt estimation θ1. The microscope head should be rotated in the direction of the minimum longitudinal position. Optimal adjustment of the microscope head is achieved when the beam along the γ= 0°-180° plane is reflected in the same vertical position in the target.
6. If the rotation angle α measured in step 3 was not appropriately adjusted, a residual tilt might be observed. Such tilt is evidenced by a misalignment (d_2_) of the reflected beam along the plane perpendicular to the microscope plane (rotation angle γ= 90°-270°). This residual tilt is adjusted by rotating the animal along the table plane by an angle of: α’= tan^-1^(tan(β)/tan(θ’)) where β is the current angle of the microscope head and θ’ is the residual tilt along the orthogonal plane, obtained with Eq. 1 using the longitudinal amplitude d_2_ measured along the γ= 90° −270°.
7. Steps 4 to 6 are repeated until the spot on the target remains at the same position on the target throughout an entire rotation of the alignment-device.

In our experience, precise alignment (see Fig. 3E) is achieved after 1 or 2 iterations. Steps 4 and 5 of the protocol are illustrated in Fig. 3D. A first rotation of the device around the objective (wherein the angle θ is color coded) reveals that the minimum position of the spot on the target is obtained for an angle γ= 30°. As such, to compensate for this mismatch, the animal has to be rotated by an angle α= −30° along the plane of the table. Rotating the device once more around the objective entails a similar spot pattern (identical amplitude), however the minimum position should be reached for γ= 0° (not shown). In this configuration, the difference between the maximal and minimum longitudinal positions (for γ= 0° and 180°) is approximately 14 mm, which can be converted to a tilt angle θ of approximately 3° (using *L* = 60 mm). For illustration purposes, rotating the microscope frontend by β= 2°, as shown in the partial tilt correction step, reduces the amplitude of the ellipse on the target. As the angle α of the sample was correctly adjusted, the minimum position is found for γ= 0°. The remaining ellipse’s longitudinal amplitude is around 4 mm, leading to a remaining tilt angle of 1°. A second correction of the microscope head by β= 1° fully compensates for the sample initial tilt θ of 3°, and the beam spot remains almost at the same position for each rotation angle γ.

**Figure 3.**
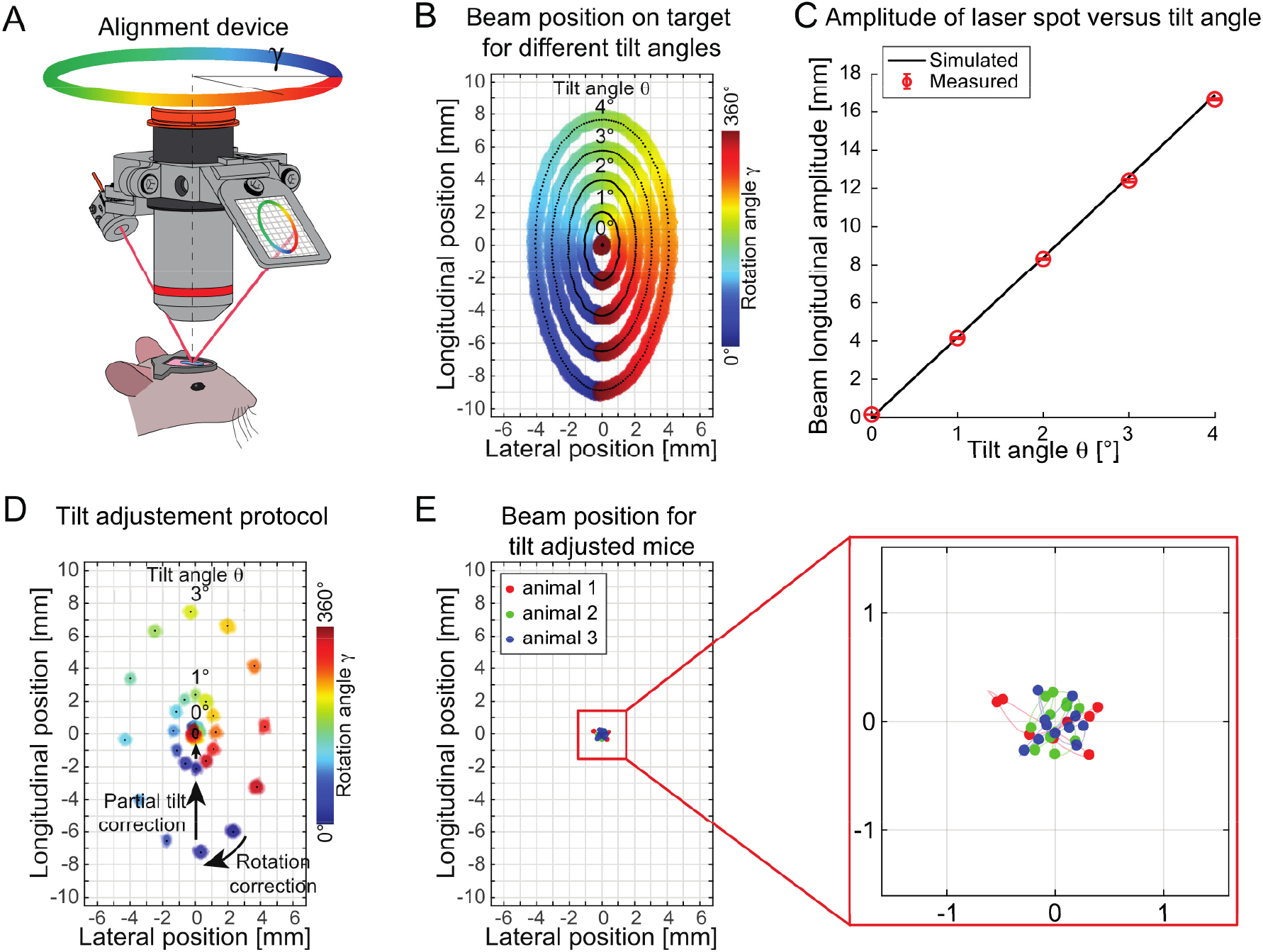
Device characterization. (A) If the glass window is tilted, rotating the device will cause the beam to wander on the target creating an ellipsoid (rotation angle of the device γ is color encoded). (B) The amplitude of the ellipsoid in the longitudinal axis is related to the tilt angle θ as predicted by the geometrical model. (C) The measured amplitudes at specific tilt angles are shown by red circles and are in good agreement with our model, shown by the black curve. (D) The tilt adjustment protocol can be summarized into two main steps; firstly, the specimen has to be rotated (α) such that the rotation angle of the alignment device (γ comprising the maximum and minimum amplitude are aligned parallel to the microscope axis (Rotation correction arrow). In a second step, the microscope head is rotated by an angle β (Partial tilt correction arrow), obtained by measuring the longitudinal amplitude of the ellipsoid and converting it into a tilt angle through Eq. 1. Ultimately, once the microscope head is set such that the tilt angle is fully compensated, the beam remains at the same position throughout an entire rotation of the device. (E) The full tilt alignment procedure was performed on 3 animals, and the amplitude of the beam on the target for a full rotation of the device was smaller than 0.5 mm on the target. The dots in panel E represent the centroids of the beam at different rotational angles and for different animals (color indicates the animal).

#### 3.2.4 Device performance

Lastly, we characterized the performance of the system in an *in vivo* setting to estimate the residual angle present after a tilt-adjustment operation. For n = 3 mice, we therefore performed the protocol described above, and captured the position of the beam on the target for different azimuth angles γ. The positions of the beam for different angles are shown in Fig. 3E, displaying a beam amplitude smaller than 0.5 mm for a full rotation. From these measurements, we calculated a residual angle of 0.15°± 0.03°, demonstrating the high sensitivity of the device.

## 4 Discussion and conclusion

Although TPEF is a powerful microscopy technique, obtaining optimal imaging performances in *in vivo* conditions can be cumbersome, as some parameters, such as the position of the animal’s imaging window are difficult to control. In this manuscript, we characterized the effects of a tilt angle between the imaging plane and the cranial window on the quality of TPEF microscopy data when imaging through a varying thickness of glass and using a high NA objective with an adjustable correction collar. In a first step, we confirmed our previous findings indicating that such a tilt angle has a deleterious effect on the imaging quality [17] and then demonstrated that this effect is further exacerbated when increasing the thickness of the glass window. Although minimizing the window thickness would be advantageous in terms of optical aberrations, using a double cover glass configuration for *in vivo* chronic brain imaging applications can be preferable as it can reduce imaging artefacts caused by brain motion.

In addition, multiple layered glass windows are not only more robust than single thinner windows, they can also limit bone growth and are therefore better suited for long-term imaging studies. Ultimately, under the experimental conditions used in this study (i.e. doubling the amount of glass), the additional layers of glass increased the impact of a phenomenon already present for single layer glass windows. As emphasized by our data, optimal imaging performance and diffraction-limited resolutions can be retrieved in both configurations by carefully positioning the sample and aligning the glass window with the imaging plane (i.e. orthogonally to the optical axis). As described in the introduction, a number of strategies have been implemented to either align the window or altogether optically cancel the aberrations caused by this tilt. Nevertheless, these techniques can be either cumbersome to implement, imprecise or costly. In view of this, we developed an affordable optical alignment device allowing estimating the tilt angle and we provide a simple procedure for its minimization. Directing a laser beam to the surface of the cranial window at different azimuth angles allows the angle mismatch between the cranial window and the imaging plane to be probed (Fig. 2 and 3) by characterizing the trajectory of the reflected beam on a target. We developed a simple model relating the observed longitudinal amplitude of the beam on the target with the tilt angle of the sample.

Although the model relies on a certain number of assumptions (e.g. the target is perpendicular to the reflected beam and the beam impinging on the surface on the rotation axis of the device), it showed a good agreement with experimental data and with a full geometrical model (not shown here). In a practical environment, due to the restricted dimensions of the glass window, we found that minimizing the trajectory of the beam on the cover glass facilitated the estimation of the amplitude of the beam on the target. The most important approximation made in this study is the assumption that the plane around which the device rotates is perfectly parallel to the imaging plane. Although this is certainly not the case in reality, in our investigations, we found that optimal imaging performance was obtained when the tilt angle was compensated. A small mismatch might therefore still be present but would be inferior to the sensitivity of the device. Ultimately, using this protocol, we were able to reduce the tilt angle in an *in vivo* setting down to < 0.2°.

An advantage of this device is that it can be easily assembled from off-the-shelf electronic and optomechanical components and 3D printed parts (the detailed drawings and a list of all required parts are available on the following link: https://huberlab.org/resources/optical-alignment-device) and does not require any additional sample preparation. It can be mounted either on the main imaging objective or on a low magnification objective with a large working distance, typically used to initially position the animal and inspect the cranial window before performing high resolution imaging.

The device is based on a simple reflection principle (i.e. equal incident and reflected angles compared to the normal of the window) and geometrical properties, which are not affected by the distance to the target, incidence point or incidence angle. This robustness makes its implementation straightforward, without any specific calibration or advanced knowledge of optics and can be used by anyone with access to a TPEF microscope. The simplicity and accessibility of the presented device are in contrast to other methods for tilt correction such as adaptive optics or iterative wavefront sensing and sample repositioning which require expensive optical components and a specialized expertise. The provided standardized protocol does not suffer from subjective visual estimations of the cranial window alignment.

Although the procedure is described here for a TPEF system possessing a rotatable frontend, it can be adapted to any tilting methods. For example, if animals are imaged in an anaesthetized state and their heads can be rotated through a tilting stage, the same strategy can be adapted to determine the head angle minimizing the tilt between the imaging plane and the cover glass.

Ultimately, the device was presented in the context of *in vivo* TPEF microscopy; however it can be mounted and used with other imaging methods (intrinsic signal imaging [20], laser speckle contrast imaging [21], third harmonic generation microscopy [22], optical coherence tomography [23]) in which the cranial window has to be aligned parallel to the imaging plane.

## 5 Acknowledgements

We thank Mario Prsa for the comments on the manuscript. This work was supported by the Swiss National Science Foundation, the European Research Council, the New York Stem Cell Foundation, Howard Hughes Medical Institute, and National Institute of Health. D.H. is a New York Stem Cell Foundation-Robertson Investigator.

